# Genomic survey maps differences in the molecular complement of vesicle formation machinery between *Giardia intestinalis* assemblages

**DOI:** 10.1101/2023.05.01.538903

**Authors:** Shweta V. Pipaliya, Joel B. Dacks, Matthew A. Croxen

**Affiliations:** Division of Infectious Diseases, Department of Medicine, University of Alberta, Edmonton, Canada; Institute of Parasitology, Biology Centre, Czech Academy of Sciences, České Budějovice (Budweis), Czech Republic; Division of Diagnostic and Applied Microbiology, Department of Lab Medicine and Pathology, University of Alberta, Edmonton, Canada; Li Ka Shing Institute of Virology, University of Alberta, Edmonton, Alberta, Canada; Women and Children’s Health Research Institute, University of Alberta, Edmonton, Alberta, Canada; Alberta Precision Laboratories, Alberta Public Health Laboratory, Edmonton, Alberta, Canada

**Keywords:** *Giardia*, Membrane Trafficking, Genome Assembly, Comparative Genomics, Population-level analysis

## Abstract

*Giardia intestinalis* is a globally important microbial pathogen with considerable public health, agricultural, and economic burden. Genome sequencing and comparative analyses have elucidated *Giardia intestinalis* to be a taxonomically diverse species consisting of at least eight different sub-types (assemblages A-H) that can infect a great variety of animal hosts, including humans. The best studied of these are assemblages A and B which have a broad host range and have zoonotic transmissibility towards humans where clinical Giardiasis can range from asymptomatic to diarrheal disease. Epidemiological surveys as well as previous molecular investigations have pointed towards critical genomic level differences within numerous molecular pathways and families of parasite virulence factors within assemblage A and B isolates.

In this study, we explored the necessary machinery for the formation of vesicles and cargo transport in 89 Canadian isolates of assemblage A and B *Giardia intestinalis*. There is considerable variability within the molecular complement of the endolysosomal ESCRT protein machinery, adaptor coat protein complexes, and ARF regulatory system. We report inter-assemblage, but no intra-assemblage variation within the trafficking systems examined. These include losses of subunits belonging to the ESCRTIII as well as novel lineage specific duplications in components of the COPII machinery, ARF1, and ARFGEF families (BIG and CYTH). Since assemblages A and B show differences in disease manifestation, our findings may well have clinical implications and even taxonomic, as the membrane trafficking system underpin parasite survival, pathogenesis, and propagation.

## INTRODUCTION

*Giardia intestinalis* is a protist pathogen responsible for the globally occurring water and food-borne diarrheal disease, Giardiasis. The World Health Organization estimates roughly 280 million cases, with greater incidence of disease occurring in resource poor regions with limited access to clean drinking water and sanitation infrastructure (Esch & Petersen, 2013). Ingestion of *Giardia* cysts contaminating food and water leads to establishment of gut infection. Chronic and recurring infections in children have developmental consequences and can extend beyond the acute intestinal disease.

Although morphologically identical, multi-locus sequence genotyping has sub-divided *Giardia intestinalis* into eight distinct assemblages (*i.e.,* genotypes), A through H, with broad animal host tropism, including humans (Heyworth, 2016). Of these, A and B have the greatest host range and are the predominant assemblages that infect humans to cause human giardiasis. Despite similarities in their ability to cause human disease, phylogenetic analyses place assemblage A closely related to the cattle-infecting assemblage E and cat assemblage F. Meanwhile, assemblage B is closer in relation to assemblage G, which infects murine hosts such as mice and rats (Cacciò et al., 2008). This suggests that zoonotic transmission into humans has occurred twice and through independent convergence.

Advancements in genome sequencing technologies have resulted in an abundance of new genomic data from numerous human and animal-infecting *Giardia* isolates for comparative analyses. Of the human-infecting isolates, assemblage A, isolate WB was the first available genome and lent tremendous insights into molecular-level losses and sequence divergence in this parasite compared to other eukaryotes (Morrison et al., 2007). Since then, other assemblage A isolates, such as AS175 and AS98, as well as assemblage B isolates, GS, GS_B, and BAH15c1, have also been sequenced and made available (Adam et al., 2013; Franzén et al., 2009; Wielinga et al., 2015; Xu et al., 2020). Recent advances in long-read sequencing technologies have also allowed cost-efficient improvements of these previous assemblies (Pollo et al., 2020; Xu et al., 2020). Other animal-infecting isolates from dog assemblages, C and D, and cattle assemblage E have also been sequenced and assembled, which have shed light on the molecular differences between the various *Giardia* assemblages (Jerlström-Hultqvist et al., 2010; Kooyman et al., 2019). Investigating these additional animal isolates is necessary due to their implications on potential for anthropozoonotic transmission between humans and domesticated animals (Fantinatti et al., 2016).

Fundamental biological and genetic differences have been enumerated between the two human-infecting strains. From an *in vitro* standpoint, compared to assemblage A, assemblage B isolates are slow-growing in axenic cultures and have lower transfection efficiency with episomally or stably integrated plasmids (Singer et al., 1998). On the other hand, assemblage A isolates are challenging to study in mice models where infection cannot be established or is cleared immediately compared to assemblage B isolates, which can readily infect and cause disease (Byrd et al., 1994). Although structurally otherwise identical, various cytogenetic differences are also present, such as the number of chromosomes per nuclei and their sizes (Adam, 1992; Tůmová et al., 2007). Although generally lower compared to other polyploid eukaryotes, genetic allelic heterozygosity (ASH) comparisons between assemblages A and B have shown variances in the levels between the two. Assemblage A isolate WB is markedly lower in overall ASH (<0.01%) compared to assemblage B isolate, GS, which has a much higher genomic allelic divergence (0.5%) (Ankarklev et al., 2010).

From a clinical standpoint, differences in human Giardiasis outcomes are also variable. Several cross-sectional clinical studies have been conducted in numerous countries (*e.g.,* United Kingdom, Scotland, Sweden, United States, Saudi Arabia, Egypt) to understand if specific strains are attributable to a higher probability of developing symptomatic or chronic disease outcomes (Cacciò & Ryan, 2008; Ferguson et al., 2020; Minetti et al., 2015a; Minetti et al., 2015b; El Basha et al., 2016). These surveys predominantly associated that *ca.* 60-65% of the infections were caused by assemblage B parasites, while the remaining 30-35% of the cases were caused by assemblage A isolates. Additionally, the latter had a greater correlation with asymptomatic or milder disease outcomes, whereas infections with assemblage B isolates had a higher likelihood of causing a prolonged or refractory disease characterized by severe diarrheal and other gastroenteritis symptoms (Lebbad et al., 2011). Patients infected with assemblage B were also less likely to respond to first-line treatments, metronidazole and albendazole (Lalle & Hanevik, 2018; Lecová et al., 2018).

Molecular-level variances within parasite biology can contribute towards these clinical disease and parasite biological differences. The increased availability of whole-genome sequencing data from numerous assemblage A and B isolates previously enabled large-scale and multi-family comparative genomic investigation and has brought forward growing evidence that suggests considerable inter-assemblage variability at the genetic level. Much of this has been found within multiple *Giardia*-specific virulence gene families, such as the cysteine-rich variant-specific surface proteins (VSPs) expressed by the trophozoites during a gut infection for parasite antigenic variation in order to modulate and evade the host immune cells (Prucca & Lujan, 2009). For example, isolates belonging to assemblage B encode almost 700 different VSPs compared to their assemblage A counterparts, which have considerably lower VSP repertoires (*i.e.,* 190 in WB isolate and 250 in the DH isolate) (Ankarklev et al., 2010; Jerlström-Hultqvist et al., 2010).

Differences in the other encoded protein machinery, particularly those belonging to the membrane trafficking system pathways, are also evident. Previous comparative genomics and phylogenetic investigations performed by us and others showed the complement and evolution of the ESCRT proteins, vesicle coats, and the ARF regulatory system proteins. Within each of these systems, consistent patterns of inter-assemblage variabilities were identified that would have substantial implications on the molecular intricacies underpinning the processes occurring at the parasite endomembrane organelles. Given the small number of isolates from each of the two human-infecting assemblages, a deeper investigation at a population level is necessary to confirm the initial findings of intra-assemblage differences and to assess what variability exists, if any within assemblages, *i.e*. at the intra-assemblage level.

Large-scale genomic data from assemblage A and B isolates are therefore essential to evaluate these trends. Recently, the British Columbia Centre for Disease Control Public Health Laboratory (BCCDC PHL) sequenced 89 *Giardia intestinalis* isolates belonging to either assemblage A or B (Tsui et al., 2018). Archived isolates were collected from fecal samples, surface waters, and beavers during periods of sporadic and regional Giardiasis outbreaks in across the province of British Columbia (Canada) between the years 1989 and 1995 as well as some from Hamilton, Ontario (Canada) (Prystajecky et al., 2015). A combination of PCR and whole-genome sequencing classified these outbreak-associated *Giardia* isolates within assemblage AI, AII, and B (Prystajecky et al., 2015; Tsui et al., 2018). In this study we leveraged the short-read sequencing data generated by the BCCDC PHL (Tsui et al., 2018) by first re-assembling the *Giardia* genomes using a ploidy-aware assembly methods, followed by a population-scale survey of the molecular complement with three important families of vesicle formation machinery, the ESCRT complexes, vesicle coats and adaptor proteins, and the ARF regulatory system proteins in these large collection of *Giardia* isolates.

## MATERIALS AND METHODS

### Paired-end read information and data retrieval

*De novo* genome assembly analyses with the BCCDC PHL isolates of *Giardia intestinalis* assemblage A and B were performed using the previously sequenced and archived paired-end inserts that were generated using the Illumina MiSeq. Cysts belonging to each isolate had been previously obtained from surface water, beaver, and human fecal samples and archived by the BCCDC PHL (Prystajecky et al., 2015; Tsui et al., 2018). Detailed geographical sources for isolate retrieval and assemblage classification were provided through Table 1 and Table S1 in the previous study by Tsui et al (Tsui et al., 2018).

Raw sequencing reads are available at National Centre for Biotechnology Information (NCBI) Short Read Archive under the BioProject ID PRJNA280606. We downloaded the sequencing data from the European Nucleotide Archive (The European Molecular Biology Laboratory, Heidelberg, Germany). Metadata detailing sequencing run statistics and biosample accessions for the 89 isolates is provided through the Supplementary Table 1 as well as in the previous study (Tsui et al., 2018).

### Read quality assessment, taxonomic classification, and de-contamination

Prior to assembly, the overall quality of the paired-end inserts was assessed using FastQC (S. Andrews, 2010). All datasets were inspected for non-*Giardia* contaminating sequences. To do so, paired-end reads belonging to each biosample were subject to taxonomic classification using the metagenomic classification software, Kraken2 (v. 2.0.8-beta), and against a custom-built database for k-mer matching against query sequences (Wood et al., 2019). The database was built by retrieving taxonomic information and full-length genomes corresponding to archaeal, bacterial, viral, human, fungal, protozoa, and vector contaminant (UniVec_Core) datasets from the NCBI Reference Sequence Database (RefSeq). In addition, genomes belonging to *Giardia muris* (GCA_006247105.1 UU_GM_1.1), *Giardia intestinalis* EP15 (GCA_000182665.1), *G. intestinalis* BAH15c1 (GCA_001543975.1), *G. intestinalis* BGS (GCA_011634595.1), *G. intestinalis* BGS_B (GCA_000498735.1), *G. intestinalis* AWB (GCA_011634545.1), *G. intestinalis* AS175 (GCA_001493575.1), *G. intestinalis* ADH (GCA_000498715.1), *G. intestinalis* assemblage C pooled cysts (GCA_902221545.1), and G*. intestinalis* assemblage D pooled cysts (GCA_902221535.1) were also added to the library using the kraken2-build option. Sequences that remained unclassified or were assigned to taxa other than the *Giardia* genus (NCBI taxid: 5741) were considered contaminants. Datasets were flagged if 20% or higher number of the total reads were not characterized as *Giardia*. These bio-samples were removed from analyses and not subject to downstream assembly process or comparative genomics. For the remainder datasets with low contamination levels, decontamination was performed using the Python script, extract_kraken_reads.py, available through KrakenTools (https://github.com/jenniferlu717/KrakenTools; Wood et al., 2019). Post-decontamination Kraken2 output was visualized using KronaTools (Ondov et al., 2011). Kraken-build shell script with all parameters has been made available as Supplementary File 1. Per-sample read taxonomic assignment is summarized in Supplementary Table 2.

### de novo genome assembly

Kraken2 analyses flagged three datasets to be highly contaminated with bacterial sequences and three with mixed assemblage taxonomic assignment. These six samples were not analyzed further. The remaining 83 paired-end filtered datasets were kept for *de novo* genome assembly using the ploidy-aware assembler MaSuRCA v. 3.3.0 (Zimin et al., 2013), using 200,000,000 hashes from Jellyfish v.3.2.4 to generate contigs and scaffolds (10-13Mb) in FASTA format (Marcias & Kingford, 2011) MaSuRCA configuration script has been made available as Supplementary File 2.

### Genome assembly completeness and contiguity evaluations

The resulting nucleotide assemblies were subject to post-assembly quality assessments by computing genome contiguity and completeness statistics. Contiguity was assessed using the N50 metric with the Quality Assessment Tool for Genome Assemblies (QUAST) v. 5.0.2, which also determined other contig statistics such as total length in base pairs and the final percent GC for each assembly (Gurevich et al., 2013). This detailed report is made available through Supplementary Table 3.

In addition to genome contiguity, to assess gene set completeness and report on the number of evolutionarily conserved near-universal single-copy orthologs, Benchmarking Universal Single-Copy Orthologs (BUSCO) v. 4.1.1 was used as a tool for a translated Hidden Markov Model-based searching against OrthoDB v.10 (Kriventseva et al., 2019; Simão et al., 2015). Run parameters for BUSCO consisted of the following: lineage dataset selection was eukaryote_odb10, analysis mode set to ‘genome,’ and TBLASTN e-value cut-off was kept as the default 1x10^-3^. All other BUSCO parameters, including those for the in-built AUGUSTUS v. 3.2.3 gene prediction software, were kept to default. All BUSCO output scores were plotted and visualized using the ggplot2 library available through the R Tidyverse package (Wickham et al., 2019). Data visualization specifications were provided through an R script (R Core Team, 2020), which is available as Supplementary File 3.

### Reference-mapped gene predictions and functional annotation

The nucleotide contigs from all isolates were used to predict gene features and annotations to screen for protein orthologs. To do so, Liftoff v.1.5.1 was used for annotation mapping from closely related reference genomes (Shumate & Salzberg, 2021).

Of the currently available assemblage A and B genomes, those belonging to *Giardia intestinalis* assemblage A, isolate WB (UU_20 assembly, GCA_011634545.1) and assemblage B, isolate BGS (GL50801 assembly, GCA_011634595.1) have been richly annotated and publicly available (Adam et al., 2013; Xu et al., 2020). Therefore, UU_20 and GL50801 assemblies were used as references for gene prediction in the A and B assemblage genomes, respectively. Input and reference genomes were first indexed and aligned using Minimap2 v. 2.17 using the following parameters: -a -eqx –end-bonus 5 -N 50 -p 0.5 (Li, 2018). Genes were successfully aligned if the coverage along the coding sequence (CDS) met a threshold of 50% or higher in sequence similarity. They were used to generate output GFF files detailing coordinates, accessions of the mapped ORFs, and accompanying gene features. Unmapped gene accessions were parsed out in a separate file. Program usage parameters were specified as per Liftoff instructions to generate corresponding GFF files and the per-sample Liftoff mapped and unmapped summaries have been made available through Supplementary Table 4.

Genome output files were used for presence/absence analyses of the vesicle formation machinery in isolates of *Giardia* assemblage A and B, previously curated through comprehensive comparative genomics and phylogenetics in preceding investigations (Leung et al., 2008; Marti et al., 2003; Pipaliya, Santos, et al., 2021; Pipaliya et al., 2021). Specifically, WB and GS accessions corresponding to subunits of the giardial ESCRT machinery, adaptins, retromer, COPII, COPI, clathrin, and the ARF regulatory system proteins, were searched into the newly generated GFF files. To consolidate these preliminary hits, assign orthology to any unmapped genes, and cross-identify and validate assemblage-specific orthologs/paralogs, TBLASTN (v. 2.2.29+) (Altschul et al., 1997) searches, with an e-value threshold set to 0.01, were also performed. Orthologs of the identified vesicle formation machinery from *Giardia intestinalis* ADH, AWB, BGS, and BGS_B were used as queries for homology searching into the contigs and scaffolds of all newly assembled genomes. Results of this survey are summarized as tile-plots, but specific scaffold locations from the TBLASTN hits have been made available through Supplementary Table 5.

Using these identified ORFs, we extracted the nucleotide sequences and performed their translation using Bedtools getfasta (Supplementary File 4). Both nucleotide as well as translated amino acid coding regions from each trafficking protein analyzed are summarized in Supplementary Table 6.

### Data availability

Final nucleotide contig and scaffold sequences for each genome have been made available through the Figshare repository (https://doi.org/10.6084/m9.figshare.12668813). For each resulting assembled genome, GFF and unmapped features files have also been made available through the Figshare repository (https://doi.org/10.6084/m9.figshare.13567655.v2).

## RESULTS

### Genome contiguity and completeness analyses yield uniform results across all isolates

Previously, Tsui and colleagues performed genome sequencing and assembly with 89 *Giardia intestinalis* assemblage A and B isolates archived at the BCCDC PHL to evaluate specific disease transmission dynamics underpinning the 1980s Giardiasis outbreak in British Columbia, with some of the isolates originating from New Zealand and Ontario to be used as reference. (Tsui et al., 2018). Although the sequenced short paired-end reads from this analysis are publicly available, the assembled genomes are not. Therefore, it was necessary to perform *de novo* genome assembly prior to comparative genomic analyses with the vesicle formation machinery, as per the summarized workflow (Figure 1). To do so, read taxonomic classification was performed first to assess contamination and remove sequences that did not correspond to *Giardia* (Figure 1). Of the 89 initially retrieved datasets, six were removed from final genome analyses, either due to considerable levels of bacterial contamination or if the samples were characterized as ‘mixed’ assemblages. In the latter case, according to Tsui et al., a mixed assemblage assignment was denoted if the sequencing reads mapped equally to both *Giardia* assemblage A and B. This does not imply existence of a genetically hybrid strain but is instead a by-product of gDNA contamination or its isolation from mixed cultures containing trophozoites belonging to both strains. Therefore, these data were treated as metagenomic. Because this investigation aimed to elucidate nuanced molecular-level differences between the two assemblages, samples with ‘mixed’ assemblage classification were also removed from analyses. This resulted in a final total of 83 datasets being used for downstream analyses, 42 of which were classified as assemblage A and 41 as assemblage B (Supplementary Tables 5 and 6).

**Figure 1.**
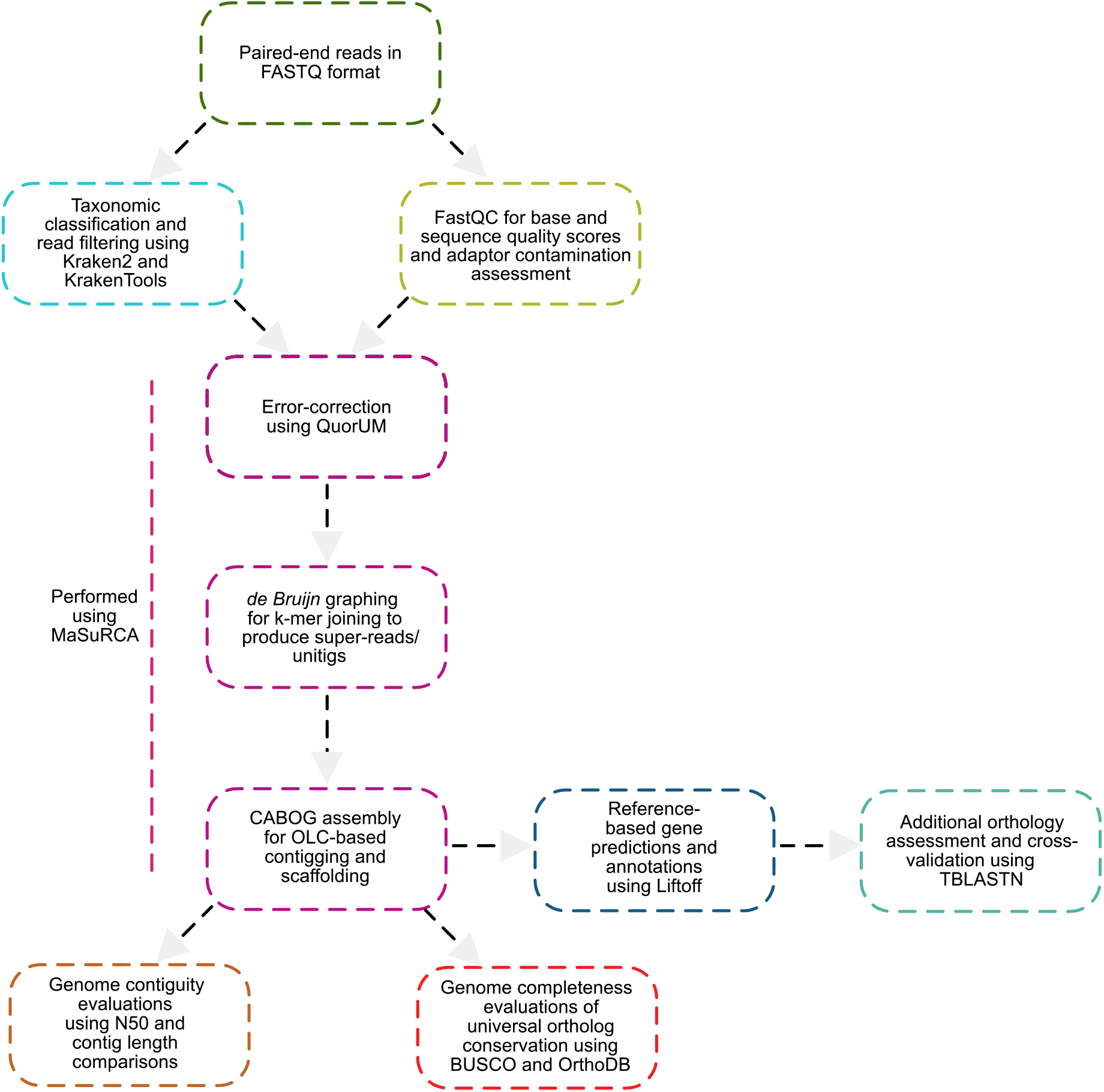
Workflow for *de novo* assembly and reference-based gene predictions. Steps included read-quality assessment and filtering using Kraken2 and FastQC, followed by *de novo* genome assembly using MaSuRCA. Assembled contigs and scaffolds were used to determine contiguity and completeness using metrics such as N50, the total number of contigs, and BUSCO scores. The resulting contigs were used for reference-based genome annotation using Liftoff for orthology mapping of genes and presence/absence analyses of the vesicle formation proteins. False negatives and existing orthology assignments were cross-validated using TBLASTN.

N50 scores and contig sizes for all assemblies were determined using QUAST. The results of this analysis calculated an average N50 of 70,543 bp and a median of 74,269 bp across all assemblies (Supplementary Tables 1 and 3). Genomes were also assembled in an average of 382 contigs that were ≥ 1kb in size (Supplementary Tables 1 and 3). In addition to genome contiguity, completeness was also assessed by determining the number of evolutionarily conserved single-copy orthologs of eukaryotic proteins by generating BUSCO scores where overall the percent BUSCO of complete plus fragmented genes was approximately 25 to 30% across all assemblies (Figure 1; Supplementary Figure 1). While these are much lower than what is typically expected of a eukaryotic genome from animal, plant, or fungal lineages (*i.e.,* 85-95%), low BUSCO scores in *Giardia* are a drawback of limited inclusion of diverse protist lineages within the eukaryote_odb10 database. In general, metamonad lineages are highly divergent in their sequences and ancestrally lack many of the proteins present in model eukaryotes, and therefore all suffer from poor BUSCO scores (Karnkowska et al., 2019; Salas-Leiva et al., 2021; Tanifuji et al., 2018). Instead of using these as absolute measures, BUSCO scores were evaluated against those published for other *Giardia* assemblies. The short-read reference genome of assemblage A, isolate WB (GCA_011634545.1) was previously determined to have a BUSCO score of C:23.9%[S:0%, D:23.9%], F:5.9%, M:70.2%, n=255, while the short-read reference genome from assemblage B, isolate GS has a BUSCO completeness score of 23.1% (https://www.uniprot.org/proteomes/UP000001548; https://www.uniprot.org/proteomes/UP000002488; Franzén et al., 2009; Morrison et al., 2007). Therefore, our re-assembled Canadian *Giardia* genomes are comparable, if not slightly more complete, in their BUSCO scores to those of the previously published short-read reference genomes belonging to isolates AWB and BGS.

### Overall GC content and genome sizes are comparable to other isolates of assemblages A and B

Aside from genome completeness and contiguity, the percent GC content (%GC) and genome size metrics for these new *Giardia* assemblies were also determined.

Of the 83 isolates assembled and kept for analyses, 42 belonged to assemblage AI or AII, and 41 to assemblage B. Previous genomic investigations with assemblage AI and AII isolates (*i.e.,* DH, WB, and AS175) were approximated to be 48% GC rich (Adam et al., 2013; Ankarklev et al., 2015; Morrison et al., 2007). A similar trend was observed across the 42 BC *G. intestinalis* assemblage A isolates, where the %GC across all isolates ranged between 47.8 and 48.9 (Figure 2B; Supplementary Table 3). A few outliers had slightly higher scores (*i.e.,* 49 to 49.96%), but overall, mean and median %GC for assemblage A were 48.5% and 48.36%, respectively (Figure 2C; Supplementary Table 3). Similar to assemblage A, previously investigated assemblage B isolates also had %GC ranging between 47 and 49 (*i.e.,* BGS, BGS_B, and BAH15c1) (Adam et al., 2013; Franzén et al., 2009; Wielinga et al., 2015). The results from this study are identical to those values, wherein the 41 BC B isolates had 47 to 49 %GC, with a mean of 48.7% and a median of 48.9% (Figure 2A and C; Supplementary Table 3).

**Figure 2.**
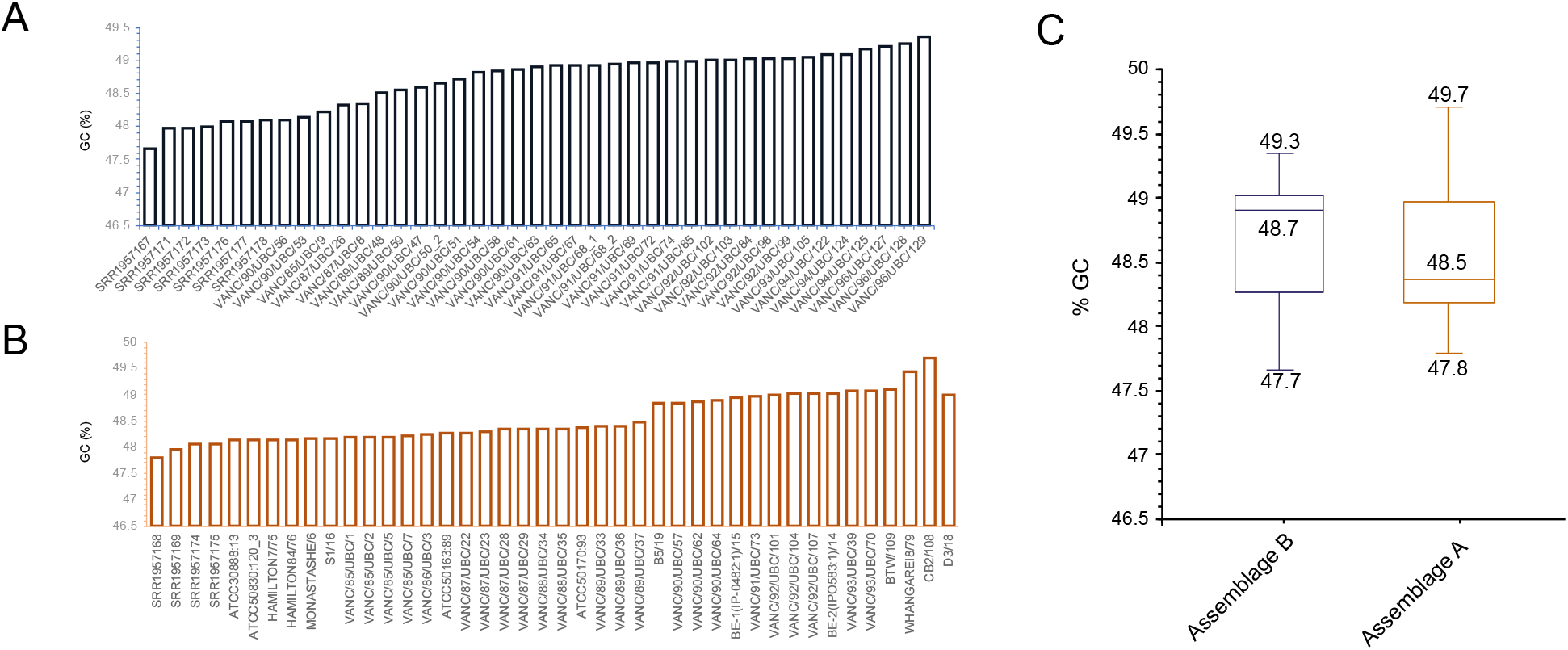
Percent GC content comparisons between assemblage A and B BCCDC PHL genome assemblies. **(A)** depicts %GC for the 41 isolates belonging to assemblage B. These ranged between a minimum of 47.7% and a maximum of 49.3%. **(B)** depicts the %GC content for assemblage A isolates, which, like assemblage B, ranged between 47.8 and 49.7. **(C)** is a box-plot depiction of the percent ranges and calculated means for GC content for assemblage A and B isolates, where a similar overall % GC was present for both (*i.e.,* ca. 48%).

Genome sizes were also evaluated and compared against the previously published *Giardia* genomes. In the literature, inter-assemblage variances within the overall genome sizes of assemblage A and assemblage B isolates have been noted, wherein assemblage AI and AII isolates (*i.e.,* WB, DH, AS175, AS98, ISS17, and ZX15) generally ranged between 10.2 Mbp and 11.7 Mbp (Adam et al., 2013; Ankarklev et al., 2015; Franzén et al., 2009; Morrison et al., 2007). In contrast, assemblage B isolates (*i.e.,* GS, GS_B, and GS/M clone H7) were comparatively larger, ranging between *ca.* 11 Mbp to 13 Mbp (Adam et al., 2013; Ankarklev et al., 2015; Franzén et al., 2009; Morrison et al., 2007). This trend was also consistent in this investigation. Apart from one outlier, the genome size range for all assemblage A isolates ranged between *ca.*10.6 Mbp and 11.9 Mbp, with an average of 11.0 Mbp and a median of 10.9 Mbp (Figure 3B and 3C). Assemblage B isolates, on the other, were larger and ranged between 10.8 Mbp and 13.7 Mbp, with an average of 12 Mbp and a median of 11.9 Mbp (Figure 3A and C). The reference isolate GS has a genome size of 12 Mbp, whereas GS_B is 13 Mbp in size. Therefore, assemblies from this study are similar in size compared to the previously published isolates from the same assemblage (Figure 3A and C).

**Figure 3.**
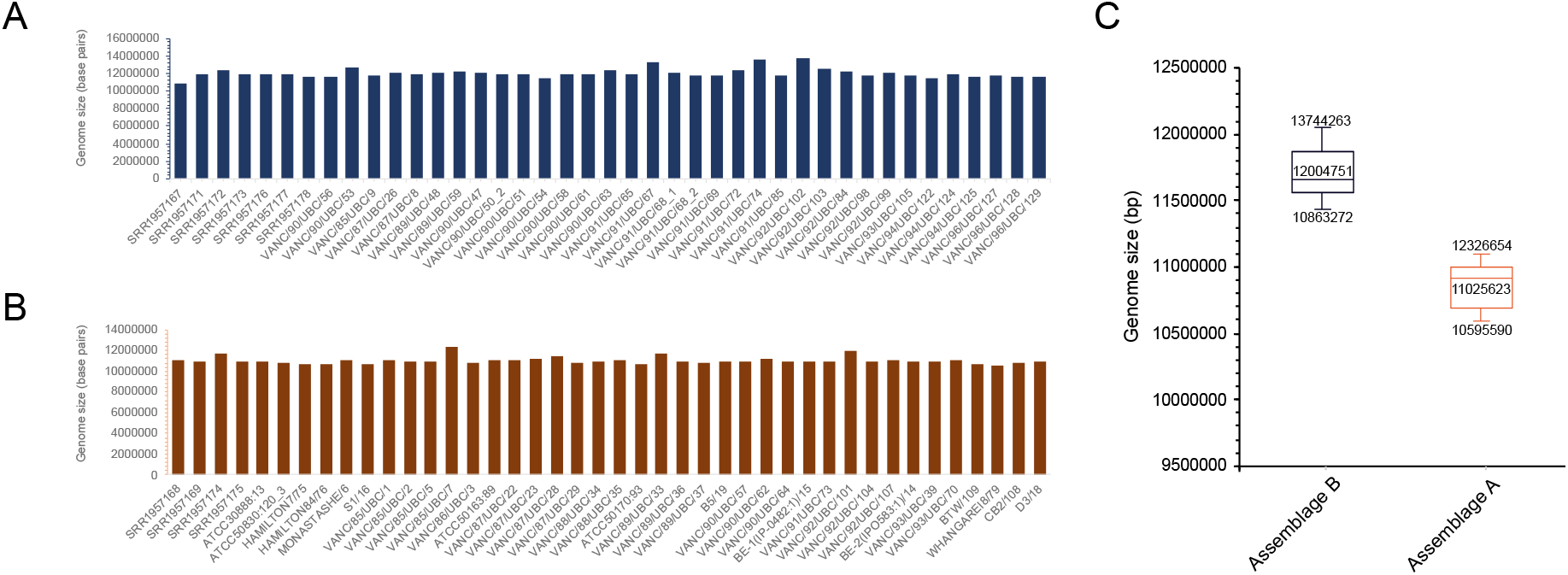
Genome size comparisons between assemblage A and B BCCDC PHL genome assemblies. **(A)** depicts genome sizes for the 41 isolates belonging to assemblage B. Genome sizes are relatively uniform across all isolates and range between 10.9 Mbp and 13.7 Mbp. **(B)** represents genome sizes for the 42 isolates belonging to assemblage A which range between 10.6 Mbp and 12.3 Mbp. **(C)** is a box-plot depiction of the range and calculated mean of the genome sizes for all isolates belonging to each assemblage. Overall, we observe assemblage A genomes to be comparatively smaller in size than assemblage B genomes.

The GC content and genome size’s collective examination yielded identical values as the previously published pan-global isolates. These findings additionally validate assembly correctness and corroborate the previously observed trends at a greater population level. Although the two assemblages are similar in their overall %GC content, isolates belonging to assemblage B are consistently larger than A.

### Gene predictions and functional annotations suggests close similarity to reference genomes

Past analyses have identified some differences in membrane-trafficking machinery complement between assemblages A and B (Marti et al., 2003; Pipaliya, Santos, et al., 2021; Pipaliya et al., 2021). However, these trends were based on a limited number of samples (three to five) per assemblage, and so a much larger sampling is needed to confirm the results. Moreover, the question of whether there are intra-assemblage differences within A or B vesicle coat machinery (Adaptins, COPI, and Retromer), COPII subcomplex, ESCRT protein complexes, and the ARF regulatory protein machinery as not been assessed at a population-level and as a result, survey of these proteins was performed with these 83 Canadian genomes to comprehend the trends previously observed in the repertoire of the endo-lysosomal vesicle formation machinery.

To do so, we first used Liftoff to map gene features and annotations from closely related species of organisms by providing the genomes from assemblage A, isolate WB (UU_20 assembly; GCA_011634545.1) and assemblage B, isolate GS (GL50801 assembly; GCA_011634595.1) as references for mapping. Many of the annotations have been curated based on experimental evidence (*i.e.,* molecular protein characterizations, transcriptomics, and proteomics) and are regularly updated by the *Giardia* research community. To supplement and cross-validate Liftoff annotations, TBLASTN analyses were also performed for genes of interest to eliminate false-negative absence artifacts due to improper or missed gene mapping.

As per recent estimations, the newly updated genome of *Giardia intestinalis* AWB (UU_20 assembly) is predicted to encode 4,963 protein-coding genes and 85 non-coding RNA (ncRNAs) genes, yielding a total of 5,048 genes (Xu et al., 2020). From this total, Liftoff mapped an average of 4,688 genes from the BC assemblage A genomes, with approximately 72 remaining unmapped (Supplementary Table 4). Similarly, albeit slightly lower, the total number of protein-coding and ncRNAs in the assemblage B reference GS genome (GL50801 assembly) is estimated at 4,470 and 92, respectively, resulting in a total of 4,562 predicted genes. An average of 4,481 genes were mapped across the BC assemblage B genomes, with 62 remaining unmapped (Supplementary Table 4). Unmapped genes may have resulted from sequence divergence or fragmented orthologs that did not share enough similarity with the target for a Minimap2 cut-off of 50%. The other possibility is that a subset of these unmapped genes are protein repertoires exclusive to the reference WB and GS genomes. Nonetheless, these findings suggest that for both assemblages A and B, approximately 98.2% of the genes and annotations from AWB and BGS genomes were successfully mapped, and therefore, encoded in the genomes of the newly assembled isolates.

### ESCRT repertoire varies between assemblage A and B isolates but is consistent within assemblages

Previously, comparative genomic and phylogenetic analyses with the late-endosomal ESCRT subcomplexes were performed by us to trace their evolution across fornicates, including several human-infecting isolates of *Giardia* (Pipaliya, Santos, et al., 2021). Here patterns of universal streamlining within ESCRTs were observed across the entire *Giardia* genus, such as the complete loss of the ESCRTI subcomplex (Pipaliya, Santos, et al., 2021). However, aside from absences, *Giardia*-specific duplication events also occurred to yield several paralogs in the following subunits: ESCRTII-Vps36 (*i.e.,* Vps36A, B, and C), ESCRTIII-Vps24 (*i.e.,* Vps24A and Vps24B), ESCRTIIIA-Vps4 (*i.e.,* Vps4A, B, and C), ESCRTIIIA-Vps46 (*i.e.,* Vps46A and B), and ESCRTIIIA-Vps31 (*i.e.,* Vps31A, B, and C) (Pipaliya, Santos, et al., 2021). Comparisons between various pan-global isolates revealed distinct differences within the repertoire of the individual subunits and the number of paralogs that comprise the giardial ESCRTII, ESCRTIII, and ESCRTIIIA. To better elucidate these inter-assemblage differences within ESCRTs and to assess if these trends exist at a population level, the repertoire of the giardial ESCRT machinery in the newly assembled BC *Giardia* genomes was reconstructed. This was done by surveying the Liftoff annotations combined with TBLASTN searches into the nucleotide contigs and scaffolds. Overall, our findings confirm and extend past results of ESCRT retention and loss within and between assemblages.

All 42 assemblage A isolates are conserved in their *Giardia* repertoire of the ESCRT complexes without any variabilities at the individual isolate level (Figure 4A). Contrary to this, previously noted streamlining exists in several ESCRT machinery components in assemblage B isolates (Figure 4B). The most prominent of these was the loss of Vps20L. The survey of 41 new assemblage B isolates confirms that this crucial ESCRTIII component is universally absent across all sampled *Giardia* assemblage B genomes (Figure 4B). TBLASTN searches using Vps20L orthologs identified in pan-global assemblage A isolates (*i.e.,* DH, WB, and AS175) were used as queries to rule out false-negative artifacts. Despite this approach, no protein hits resembling Vps20L orthology were identified. Similar trends were observed within one of the three Vps4 paralogs (*i.e.,* Vps4C), which also remained unidentified in the pan-global isolates of assemblage B. Using both Liftoff and TBLASTN, only Vps4A and Vps4B were retrieved across all 41 isolates (Figure 4B). Therefore, the absence of Vps20L and Vps4 points towards a global loss of these components in this assemblage.

**Figure 4.**
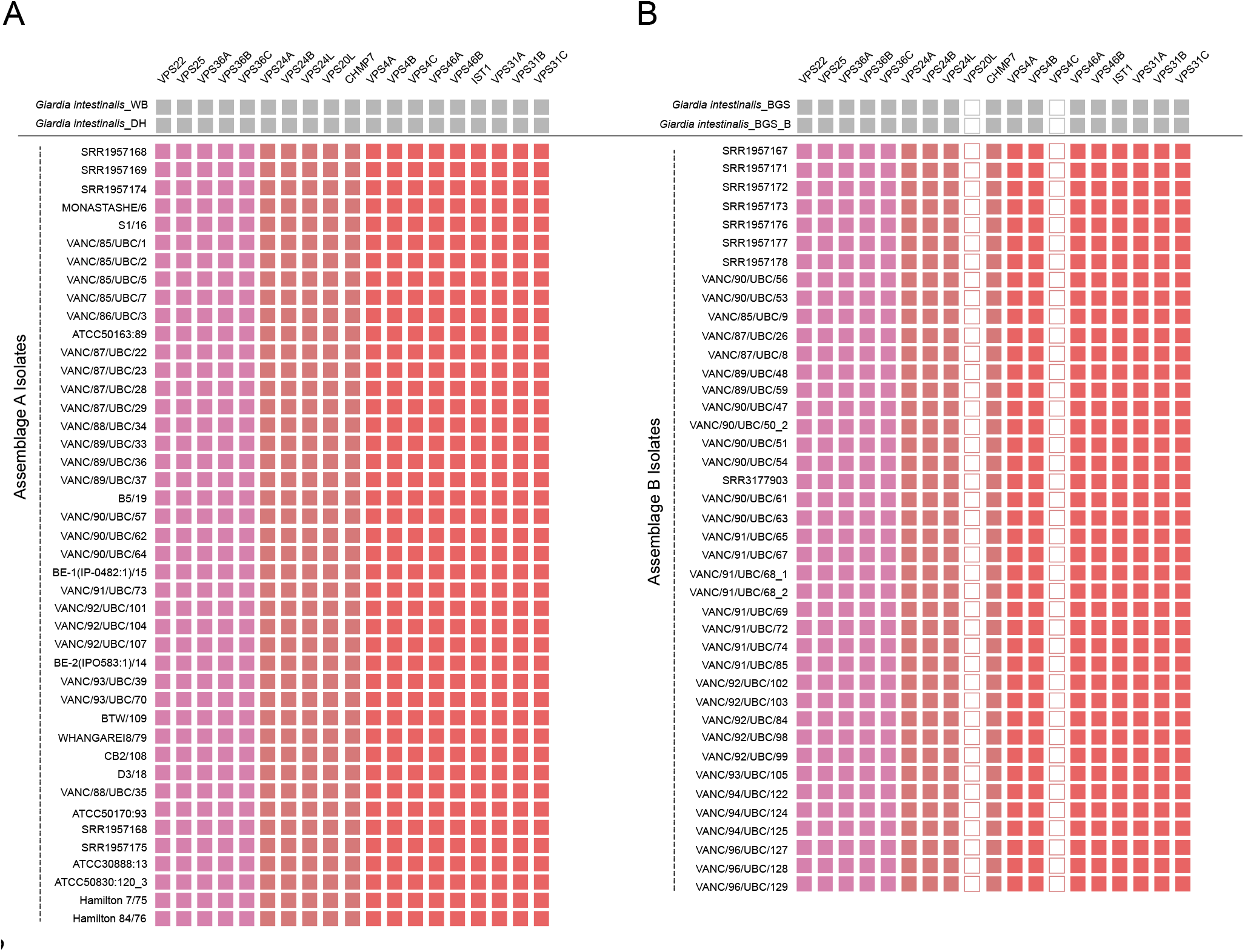
Tile-plot depictions of the ESCRT repertoire in the BCCDC PHL isolates. **(A)** depicts the giardial ESCRT repertoire distribution in the newly assembled BCCDC PHL isolates compared to the two pan-global reference assemblage A isolates, WB (AI) and DH (AII). No absences were identified within any assemblage A isolates. **(B)** shows the distribution of the giardial ESCRT repertoire in the newly assembled BCCDC PHL assemblage B isolates compared to the two pan-global referece assemblage B isolates, BGS and BGS_B. Assemblage-specific losses were identified within the ESCRTIII-Vps20L and ESCRTIII-Vps4C, and are consistent with the findings in the pan-global isolates, indicated in grey.

Altogether, these findings validate previous observations of molecular differences within the endo-lysosomal ESCRT subunits between the two assemblages and allow for an extension of those conclusions at a greater population level.

### Vesicle coat complexes follow a pattern of inter-assemblage molecular differences

Identical to the analyses performed with the ESCRTs, the evolution and the molecular complement of the heterotetrameric complexes (adaptins and COPI)(Marti et al., 2003). To consolidate those findings as well as to identify any other possible isolate-level differences, the *Giardia* complement of HTACs was determined 83 BCCDC assemblage A and B genomes were determined.

The giardial repertoire of adaptins, COPI, retromer, and clathrin heavy chain were conserved from an inter-assemblage standpoint (Figure 5). However, isolate-level variabilities were present within the adaptins.

**Figure 5.**
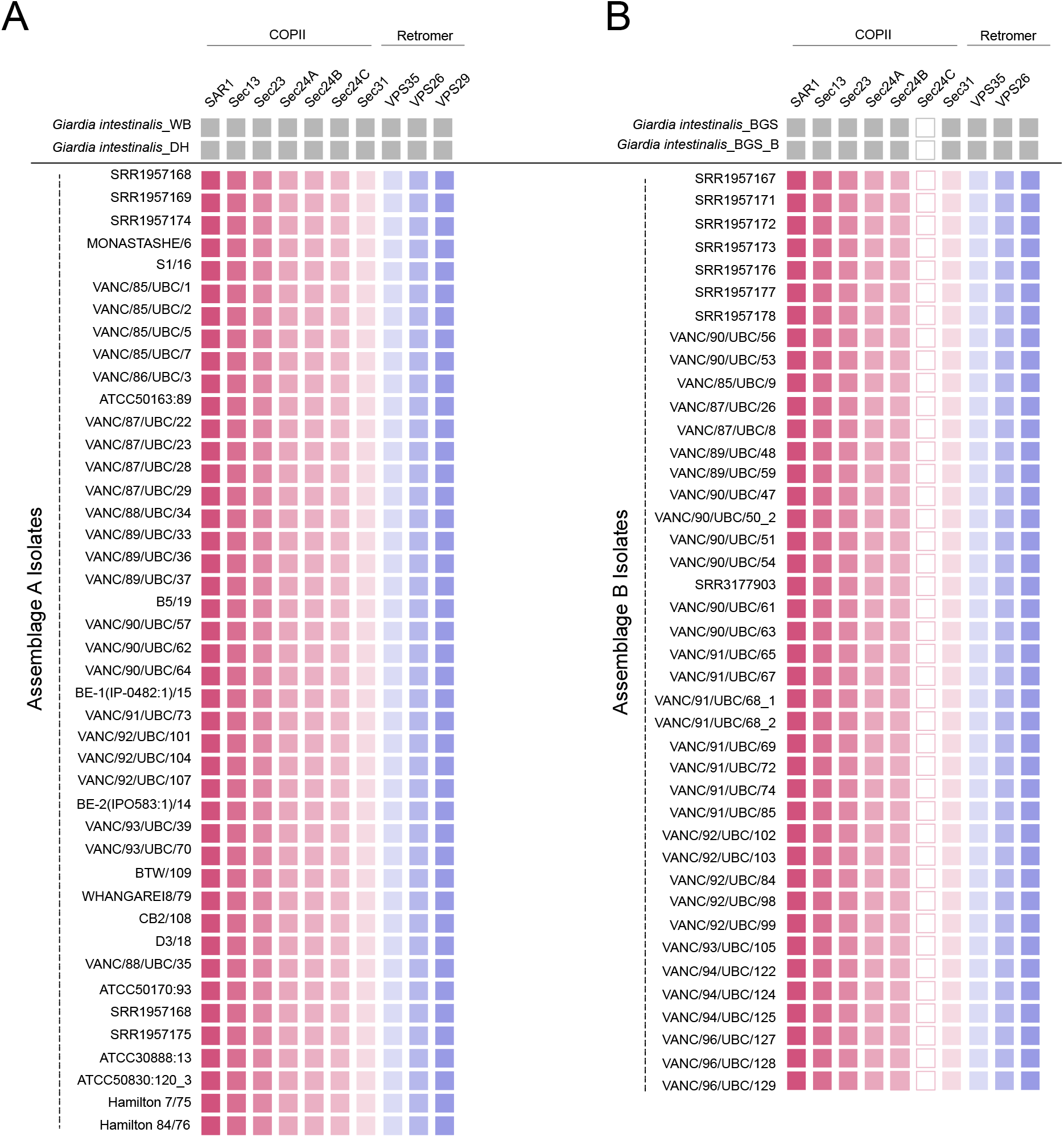
Tile-plot depictions of COPII and retromer repertoire in the BCCDC PHL isolates. **(A)** depicts the distribution of the giardial COPII and retromer components in the newly assembled genomes of the BCCDC PHL isolates, compared to the two pan-global reference assemblage A isolates, WB (AI) and DH (AII). No absences were identified within any of the assemblage A genomes. **(B)** depicts the distribution of the giardial COPII and retromer repertoire in the newly assembled BCCDC PHL assemblage B genomes and compared with the two pan-global reference assemblage B isolates, BGS and BGS_B. Assemblage B-specific losses are evident within one of the paralogs of COPII-Sec24FII (Sec24C).

Unexpectedly, two instances of gene absences within components of the AP-1 subcomplex in genomes of assemblage B were noted. First was the lack of AP-1μ in the SRR1957167 assembly and the second absence was within the AP-1σ subunit in the VANC/96/UBC/129 assembly (Figure 6B). TBLASTN searching using AP-1μ and AP-1σ orthologs from other assemblage B genomes could not identify μ1 or σ1 subunits in these genomes other than those belonging to the AP-2 sub-complex. The absence of AP-1μ and AP-1σ, although surprising, is highly unlikely and should be ascertained as a technical assembly or sequencing-related issue. AP-1μ is absent from the SRR1957167 belonging to assemblage B, which is an outlier as it has a comparatively smaller genome size (10.9 Mbp) and lower N50 (44, 795 bp) with its counterparts. Both attributes are likely a result of low starting sequencing depth compared to the other isolates and hence a cause for gene absence artifacts. In the case of AP-1σ, which is missing from the VANC/96/UBC/129 assembly, may have been due to low coverage k-mers in this region and therefore discarded during the assembly process. These are more likely scenarios than instances of actual loss, as the remaining 39 isolates from assemblage B encode both adaptin subunits.

**Figure 6.**
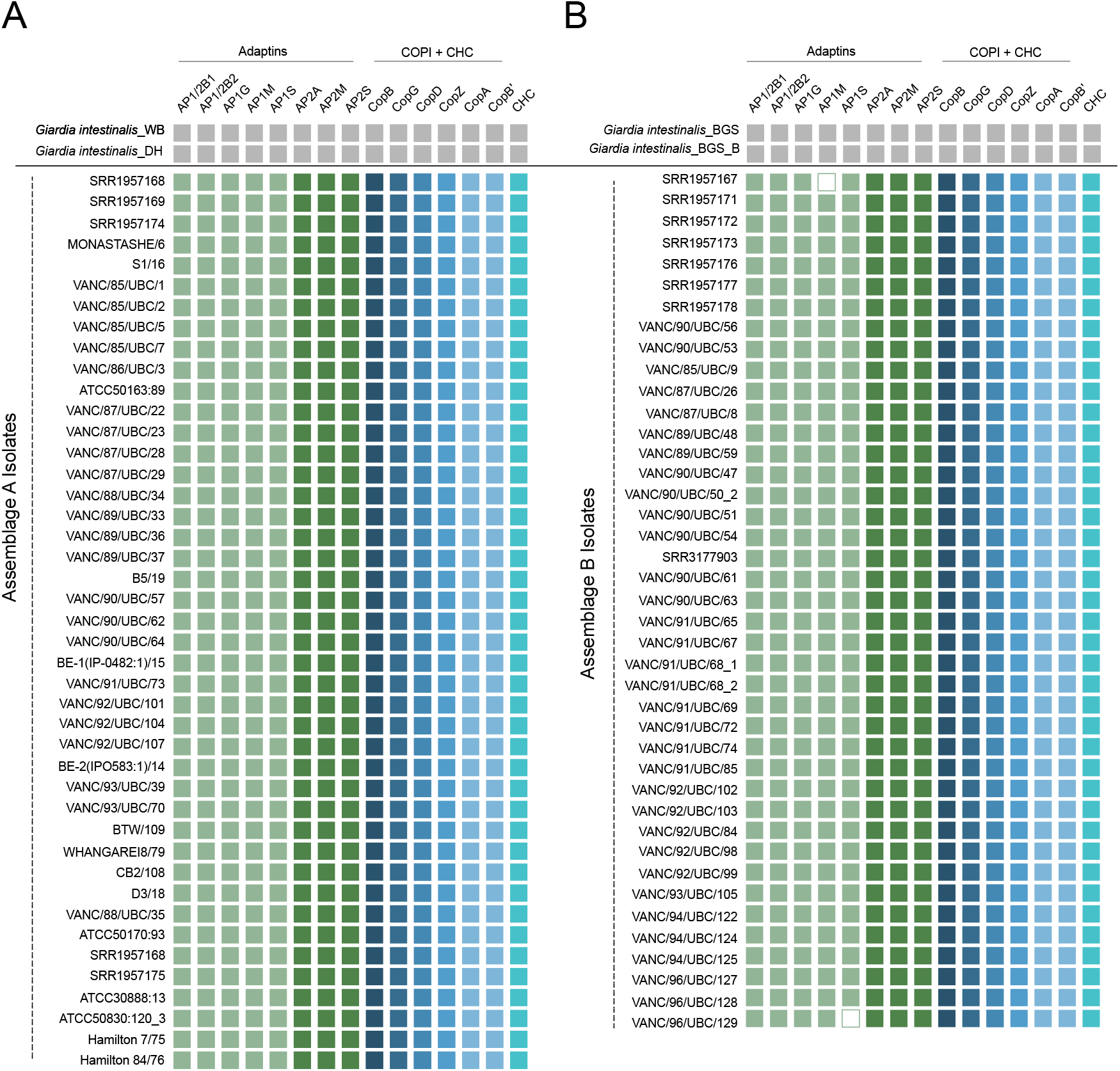
Tile-plot depictions of heterotetrameric adaptor complexes and clathrin repertoire in the BCCDC PHL isolates. **(A)** depicts the distribution of the previously identified HTACs and clathrin components in the newly assembled BCCDC PHL isolates and compared with the two pan-global reference assemblage A isolates, WB (AI) and DH (AII). No absences were identified in any assemblage A genomes. **(B)** depicts the distribution of adaptin, COPI, and clathrin components in the newly assembled BCCDC PHL assemblage B genomes in comparison to the two pan-global reference assemblage B isolates, BGS and BGS_B. Although no large assemblage-wide losses are evident, instances of individual isolate-specific absences within AP-1 subunits (*i.e.,* AP-1μ and AP-1σ) were present.

The molecular complement of COPII, retromer, and clathrin have not been examined at the same depth and as recently the above systems and therefore were examined here. For nearly all components we found the identical repertoire across Retromer, Clathrin, and most of COPII. Unexpectedly, we find multiple paralogs of the COPII component Sec24, which we have termed Sec24A, B, and C across all 41 assemblage A genomes (Figure 5). However, COPII-Sec24C was not found in any of the assemblage B genomes, despite searching by TBLASTN analyses using assemblage A Sec24C sequences (Figure 5B).

### Paralogues of the ARF regulatory system proteins continue to differ between the two assemblages

Finally, although ancestral losses shaped the fornicate ARF regulatory system (Pipaliya et al., 2021), surprisingly tight conservations within the overall complement of ARF1, ARF GAPs, and ARF GEFs exists in *Giardia* and its free-living relatives. Despite this, there remained critical differences in the encoded proteins of the two assemblages. Namely, paralogs of the fornicate-specific ARF1F proteins and Sec7 domain-containing ARF GEFs differed (Pipaliya et al., 2021).

In this previous survey, of the three fornicate-specific ARF paralogues traced in *Giardia intestinalis*, assemblage A possessed all three ARF1 variants (*i.e.,* ARF1FA, ARF1FB1, and ARF1B2), identified, whereas a secondary loss within one of the two ARF1FB paralogs (*i.e.,* ARF1FB2) was identified in the 41 assemblage B examined (Pipaliya et al., 2021). Extending this to the 83 genomes, this was confirmed, whereby all new assemblage A genomes possessed the three paralogs, while ARF1FB1 was universally absent across all 41 assemblage B genomes (Figure 7B). Unlike the previously discussed systems where absences were strictly observed in assemblage B, that was not the case with the ARF regulatory system proteins. Assemblage A has been reported to lack one of the two BIGL proteins, while assemblage B did not possess one of the two Cytohesin identified in assemblages A and E (Pipaliya et al., 2021). Once again, the population-level survey confirms the absence of one of the two BIGL proteins from assemblage A and CYTHL from assemblage B (Figure 7), but no intra-assemblage variability.

**Figure 7.**
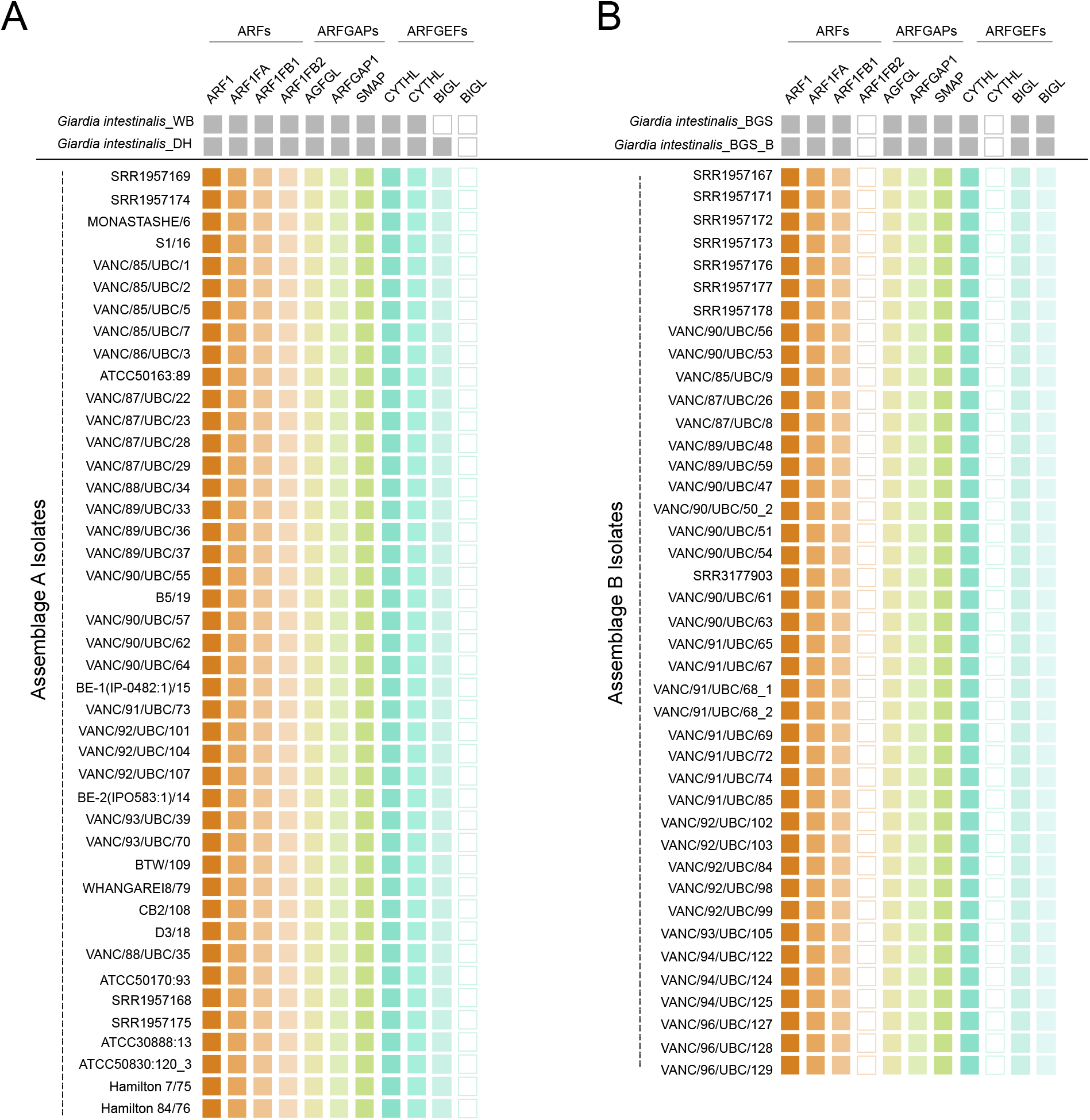
Tile-plot depictions of the giardial ARF regulatory system protein repertoire in the BCCDC PHL isolates. **(A)** depicts the distribution of the previously identified giardial ARF, ARF GEF, and ARF GAP repertoire in newly assembled BCCDC PHL *Giardia* genomes compared to the two pan-global reference assemblage A isolates, WB (AI) and DH (AII). In the previous survey, isolate WB was shown to lack both paralogues of BIG (Pipaliya et. al., 2021). However, loss within only one of the two BIGL paralogues is apparent in the new BCCDC assemblage A isolates. **(B)** depicts the distribution of the giardial ARF regulatory system proteins in the newly assembled BCCDC PHL assemblage B isolates in comparison to the two pan-global reference assemblage B isolates, BGS and BGS_B. Assemblage-wide losses within ARF1FB2 and one of the two CYTHL paralogs are also evident and consistent with the previous results.

## DISCUSSION

Past molecular evolutionary work has identified distinct differences within subunits of the trafficking complexes encoded by the two human-infecting *Giardia intestinalis* assemblages. Because the previous sampling was limited to few genomes, a population-scale investigation to better define the heterogeneity within these protein complements was necessary. The assembly of 83 full-length genomes belonging to Canadian isolates of *Giardia intestinalis* assemblage A and B permitted a large-scale comparative analysis of the vesicle formation machinery.

### Genome size differences and implications for parasite cell biology

Because it was assumed that the two human-infecting assemblages were be genetically distinct strains, an impartial examination into the differences between the trafficking complement and other genomic attributes (*i.e.,* genome size and GC content) could have only been made appropriately through a *de novo* approach. Therefore a *de novo,* instead of reference-aligned assembly, approach was pursued for genome assembly to limit reference bias and account for possible structural and sequence variants.

Using MaSuRCA, we assembled a roughly equal number of genomes belonging to both assemblage A and B isolates (*i.e.,* 42 assemblage A and 41 assemblage B genomes). We then compared the %GC and genome sizes against one another and with the previously published pan-global isolates. Currently, there exist a total of 12 publicly available assemblies for both assemblages. Upon further breakdown, approximately half belong to AI and AII, which consist of the following isolates: WB, DH, AS175, AS98, ISS17, and ZX15 (Adam et al., 2013; McArthur et al., 2000; Morrison et al., 2007; Weisz et al., 2019; Xu et al., 2020). When these isolates are examined for their %GC, a range of 48.5 to 49.5% is obtained, apart from AS98, which is markedly lower in its %GC (45.50). In comparison, %GC of the BC assemblage A genomes yielded an average of 48.5%, which is akin to the GC content of the previously published AI and AII genomes. Within assemblage B, previously published short-read assemblies consist of GS, GS_B, GS/M clone H7, and BAH15c1 isolates (Adam et al., 2013; Franzén et al., 2009; Weisz et al., 2019; Wielinga et al., 2015). Like assemblage A, the %GC for these genomes ranged between 47.0 to 49.2%. The average %GC for BC assemblage B genomes from this investigation was also comparable (48.7%). Combining these results, *Giardia intestinalis* assemblage A and B isolates are similar in their overall GC content to the values reported in the literature, and little variability exists at the inter-assemblage level.

In contrast, findings from these investigations, in combination with those from previous studies, indicate that the two assemblages vary in their overall genome sizes. The BC assemblage A isolates ranged between 10.6 to 12.3 Mbp, similar to WB, DH, AS175, AS98, ISS17, and ZX15, which also ranged between 10.3 to 12.08 Mbp (Adam et al., 2013; Morrison et al., 2007; Weisz et al., 2019). In comparison, assemblage B genomes were larger (*i.e.,* 11 to 13.7 Mbp) with similar genome sizes as GS, GS_B, GS/M clone H7, and BAH15c1 that ranged between 10.4 to 13.6 Mbp. These inter-assemblage differences in genome sizes have implications on the encoded protein-coding repertoire as well. Although the BUSCO scores suggest conservation in 27% of the eukaryotic orthologs in all genomes, previous comparative genomic differences have identified several considerable cytogenetic differences between assemblages A and B isolates. These are within structural organization of chromosomes, protein-coding syntenic regions, encoded repertoires of gene families, and inter-assemblage sequence divergences in conserved protein orthologs.

Pulsed-field gel electrophoresis for separation and comparative assessment of molecular weights of the five *Giardia* chromosomes (Chr) belonging to isolate WB and GS revealed distinct differences. WB has chromosomes that are 1.5, 2, 3, and 3.5 Mbp in size. On the other hand, GS has a slightly larger Chr1 (1.8 Mbp) due to a recombination event with Chr2 (Upcroft et al., 1996, 2010). Optical mapping between these five chromosomes identified considerable structural rearrangements, inversions, and translocations (Adam et al., 2013). Notably, even within the syntenic regions, only 70% sequence similarity was present between GS and WB. Comparisons between the number of protein-coding genes were also dissimilar, where ortholog overlap analyses identified 2962 heterologous open reading frames (ORFs) in assemblage B isolate GS_B. In contrast, assemblage A isolates, DH and WB, were reduced in this number by half, where only 1935 and 1067 unique ORFs were identified, respectively (Adam et al., 2013). Within these ORFs, the number of genes corresponding to major families of virulence genes, especially in the cysteine-rich variant surface proteins (VSPs), differed significantly. In assemblage B isolate GS_B, approximately 6.7% of all predicted open reading frames corresponded to VSPs, whereas assemblage A isolates, DH and WB, were comparably lower in their proportion of the encoded VSPs, ranging at *ca.* 3-4% (Adam et al., 2013). Additional sequence level assessments of the shared VSP profiles only yielded a maximum of 55% sequence similarity between the two assemblages (Ankarklev et al., 2015). Identical differences likely exist between isolates of assemblage A and B examined here.

### Potential impacts of missing vesicle formation machinery on secretory processes between the two assemblages

Previous functional work has determined the role of the vesicle formation machinery within *Giardia* to be critical to the parasite’s ability to facilitate secretory and uptake processes in trophozoites and encysting cells (Faso et al., 2013; Hehl et al., 2000; Miras et al., 2013; Pipaliya, Santos, et al., 2021; Rivero et al., 2012; Touz et al., 2004; Zumthor et al., 2016). Therefore, divergences in its molecular complement could implicate crucial functional variabilities within giardial trafficking processes at these organism-specific compartments.

Within ESCRTs, variations in ESCRTIII-Vps20L and ESCRTIIIA-Vps4 were identified. Cellular localization investigations by us and by previous others in the field have lent unparalleled insights into the promiscuity of ESCRTs at different giardial endomembrane compartments. *Giardia* ESCRTIII-Vps20L was primarily localized to the ER in conjunction with numerous ESCRT subunits (Pipaliya, Santos, et al., 2021). While the specific dynamics of inter-ESCRT association by sequential recruitment onto various organellar membranes are currently unknown in *Giardia*, some conserved aspects such as cargo recognition and membrane remodelling must still exist at those different compartments. Therefore, the absence of critical subunits such as Vps20L in assemblage B isolates implies possible functional compensation mechanisms to fulfill those same roles or that there is divergence in the underlying pathway wherein Vps20L is simply not necessary in this assemblage. While the former scenario may very well be the case for the missing paralogs such as Vps4C, which may be having redundant cellular roles as other Vps4 proteins at the peripheral vacuoles in assemblage A isolates, the latter scenario could signify crucial biological differences. Differences in paralog functions and molecular association of the giardial trafficking machinery, including the ESCRTs (*i.e.,* Vps4 and Vps46), have been hinted at previously and therefore is a plausible scenario, but one that still requires robust molecular and biochemical testing (Saha et al., 2018). Nonetheless, these results posit a potentially differentiated underlying mechanism by which the existing ESCRT subunits function within the *Giardia* trophozoites of the two assemblages.

Like the ESCRTs, although most vesicle coats (*i.e.,* adaptins, COPI, and retromer) are conserved in their complement between the two assemblages, differences were noted within the COPII components specifically in the number of lineage-specific paralogs of Sec24. Giardial COPII machinery is fundamental to ESV biogenesis and maturation to ensure the transport of cyst-wall material to the parasite surface (Faso et al., 2013; Stefanic et al., 2009). Sec24, along with Sec23, forms pre-budding complexes upon cyst-wall protein recognition for a COPII-coat assembly at the ER-exit sites (Faso et al., 2013; Zamponi et al., 2017). In the absence of one of the three Sec24 paralogues, it could be that the remaining two Sec24 proteins may be fulfilling this role in assemblage B isolates and that plasticity within this system exists to accommodate losses such as this one. A contrary scenario may also be possible, wherein absence in Sec24 paralogs could indicate differences in COPII-assembly dynamics. All fornicates lack Sec16, which typically stabilizes Sec23/Sec24 and Sec13/Sec31 coats during the vesicle budding process. Therefore, ancestral neofunctionalization of these lineage-specific paralogs may have occurred to compensate for missing Sec16 roles. A secondary loss of this protein in *Giardia* assemblage B could then have implications on the molecular dynamics of how the COPII-coat is stabilized in those parasites, which in turn could translate to altered ESV-biogenesis mechanisms or cyst-wall material trafficking. *In vitro* encystation has so far not been possible in assemblage B, and therefore, these postulations remain untestable conjectures (Barash et al., 2017).

Finally, differences in the ARF regulatory system proteins, especially the ARF1F GTPases, again may indicate functional redundancy of these paralogues in assemblage B at the PECs. Differences in the GEF complement may indicate variable regulation of the giardial ARFs. While testing these scenarios through fluorescent microscopy and proteomics experiments in the trophozoite stages of assemblage B isolates would be interesting, it is challenging to generate and establish parasite cultures of transgenic variants of assemblage B isolates, as they are slow growing and not amenable to stable or episomal integration of plasmids for recombination protein expression. Once technical advancements are made in this area of *Giardia* biology, these experiments should be pursued as they will open avenues to elucidate novel ARF biology and GTPase regulation mechanisms in *Giardia* and eukaryotes in general.

### Giardia as a species-complex

Most tantalizingly, our findings support previously reported genomic and molecular complement-level differences at the population level. The definition of what constitutes ‘species’ remains a hotly debated philosophical discussion within the fields of ecology, evolution, and medicine. These lines are especially blurred in microbiology. Historically, the biological species concept has defined two organisms to belong to the same species if, through sexual reproduction, they can produce viable progeny (De Queiroz, 2005). There is limited evidence for sexual reproduction or inter-assemblage genetic exchange in *Giardia*. Therefore, their species status under the strict umbrella of the biological species concept remains contested. On the other hand, the phylogenetic species concept defines species as a group of organisms with shared and unique evolutionary histories that possess a defined set of traits (De Queiroz, 2007). Phylogenetic placement using multi-locus sequence genotyping places *Giardia intestinalis* assemblages A and B as paraphyletic lineages nested within other animal-infecting assemblages (Cacciò et al., 2008; Huey et al., 2013; Lee et al., 2020). This implies that zoonoses for human pathogenesis occurred convergently. Combined with these phylogenetic classifications, differences in genome attributes and encoded protein repertoires strengthen the notion that *Giardia intestinalis* assemblage A and B (and all other assemblages) can be defined as different species as per the phylogenetic-species concept. Additionally, previous genome comparisons estimate only 77% nucleotide-level sequence conservation and 78% amino-acid similarity in the protein-coding regions between assemblages A and B isolates (Franzén et al., 2009). Genome recombination analyses performed on assemblage A, B, and E isolates further supported this notion (Xu et al., 2012). Therefore, from evolutionary, molecular, and genomic standpoints, the combined evidence further support the idea that *Giardia intestinalis* assemblages as separate species, as has been proposed by many other *Giardia* genomic and molecular biologists (R. H. Andrews et al., 1989; Cacciò et al., 2018; Seabolt et al., 2022; Wielinga et al., 2023).

Overall our findings are that there exist limited but notable differences between the two human-infecting *Giardia* assemblages, but little to no variability within the Canadian isolates that we examined. These genomic differences, and unity respectively, point to potential cell biological explanations of membrane-trafficking and ultimately clinical differences between the assemblages, and bolster the argument that these may in fact be cryptic species. Regardless, the 83 genome assemblies will provide a wealth of new genomic data for biologists to explore.

## ACKNOWLEDGMENTS

This work was supported by a CIHR Doctoral Scholarship (CIHR CGM 175863) and an Alberta Innovates Graduate Student Scholarship awarded to Shweta V. Pipaliya. Research in the Dacks lab is funded by NSERC Discovery Grants (RES0043758, RES0046091).

