## Supplementary Figure 1 for "Genomic survey maps differences in the molecular complement of vesicle formation machinery between *Giardia intestinalis* assemblages"

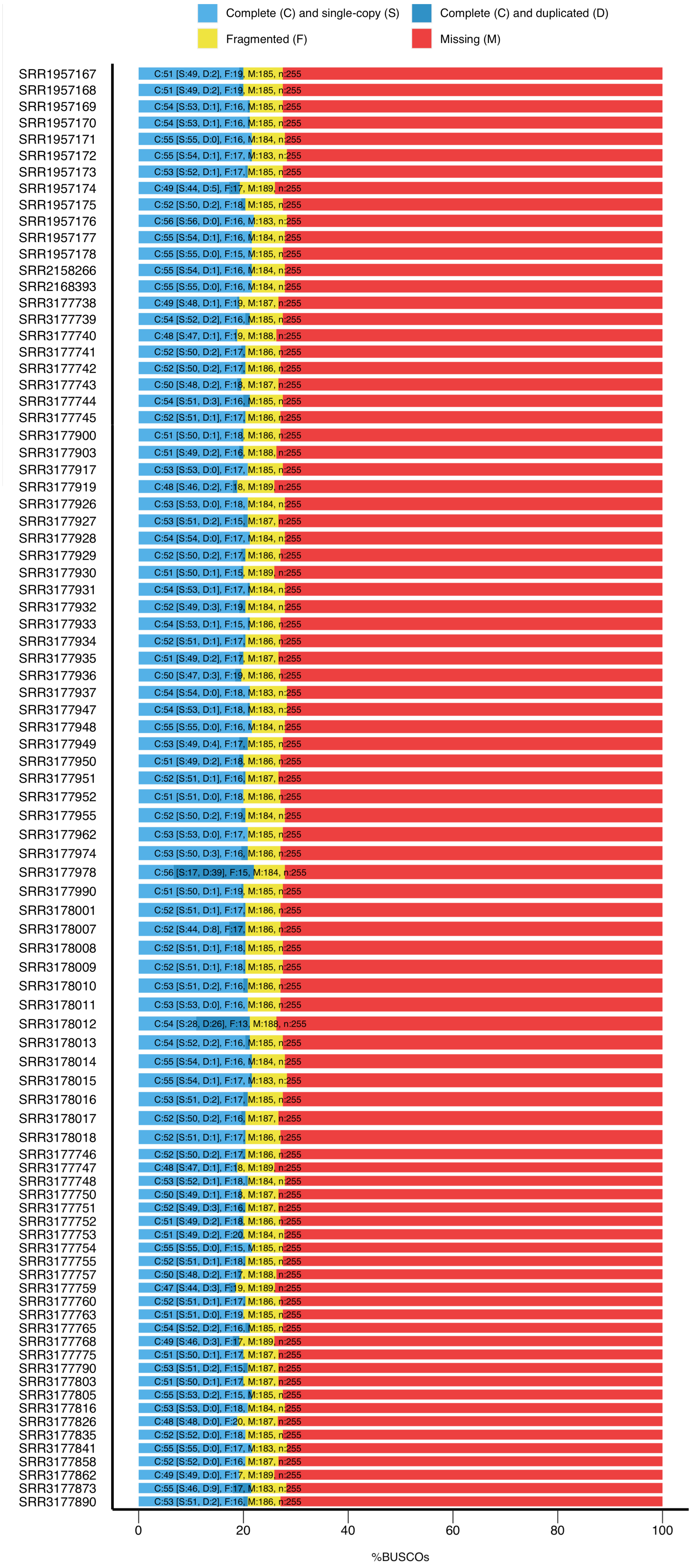


**Figure S1. BUSCO Assessment results**. Summary plot depicting BUSCO results for the 89 Canadian assemblies generated in this study. Based on the 255 single-copy orthologs from the eukaryotic ortholog database (version 10), the percent of complete and fragmented genes (blue and yellow portion of the bars) conserved across *Giardia* assemblage A and B ranged between 25 to 30% across all assemblies. Red portion of the bar represents missingness.
