## Supplementary File 4 for "Genomic survey maps differences in the molecular complement of vesicle formation machinery between *Giardia intestinalis* assemblages"

# 2022-07-12
#
# given a list of reference accession IDs, pull out the nt and aa sequence in tabular format
# list contents:
# col 1 accession number
# col 2 gene annotation
# col 3 gene class
# col 4 assemblage

# bioawk awk version 20110810
# bedtools v2.29.2
# seqkit v2.2.0

if [[ $1 == "" ]]
then
	echo "usage:" $0 "--fasta genome.fasta --gff genome.gff --list list.txt --prefix prefix"
	exit 1
fi

while [[ "$1" != "" ]]
do
	case "$1" in
		--fasta)
			shift
			fa="$1"
			prefix=$(basename "$fa")
			;;
		--gff)
			shift
			gff="$1"
			;;
		--list) 
			shift 
			lst="$1"
			;;
		--prefix)
			shift
			prefix="$1"
			;;
	esac
	shift
done

if [[ ! -f "$fa" ]]
then
	echo fasta file: "$fa" is missing.
	exit 1
fi

if [[ ! -f "$gff" ]]
then
	echo gff3 file: "$gff" is missing.
	exit 1
fi

if [[ ! -f "$lst" ]]
then
	echo list file: "$lst" is missing.
	exit 1
fi

# make a temporary directory
tmpdir=$(mktemp -d -t XXXXXXXXX)


# make it look for the accession with tag -t26_1 which seems to be applied by LiftOff (greater specificity)
# eg. GL50581_244-t26_1
# only capture and translate CDS

# once the accession is found in the GFF (column $9) (and its a $3 == CDS), use the coordinates to pull out
# the nucleotide sequence with bedtools getfasta. Then seqkit translate can translate into amino acids
# bioawk is used to wrangle some of the data

while read accession annotation class assemblage
do

	bedtools getfasta \
		-s \
		-fi "$fa" \
		-bed <(bioawk -c gff -v accession="ID=""${accession}""-t26_1" '$9 ~ accession && $3 == "CDS"' "$gff") > "${tmpdir}"/tmp.o
	
	# output should be in tmp.o in the temporary directory
	if [[ -s "${tmpdir}"/tmp.o ]]
	then
		seqkit translate --clean "${tmpdir}"/tmp.o \
			| bioawk -c fastx -v accession="${accession}" -v annotation="${annotation}" -v prefix="${prefix}" -v class="$class" -v assemblage="$assemblage" '{
				if (FNR==NR) {
					aa=$seq
				}
				else {
					# get contig name from $name eg scf7180000002853:13782-14391(+)
					split($name,contig,":")
					# get start and stop coordinates eg 13782-14391(+)
					split(contig[2],co,"(")
					split(co[1],coord,"-")
					# get strand from co[2] eg +)
					split(co[2],strand,")")
					
					print prefix,contig[1],coord[1],coord[2],strand[1],accession,assemblage,class,annotation,$seq,aa
				}
			}' /dev/stdin "${tmpdir}"/tmp.o
	fi
	

done < "$lst"

# remove the temporary directory
rm -rf "${tmpdir}"

exit 1
