## Supplementary File 2 for "Genomic survey maps differences in the molecular complement of vesicle formation machinery between *Giardia intestinalis* assemblages"

#!/usr/bin/env bash

#Illumina paired end reads supplied as <two-character prefix> <fragment mean> <fragment stdev> <forward_reads> <reverse_reads>
#if single-end, do not specify <reverse_reads>
#MUST HAVE Illumina paired end reads to use MaSuRCA

PE= pe 250 38 *_1.fastq *_2.fastq

END

PARAMETERS

#PLEASE READ all comments to essential parameters below, and set the parameters according to your project

#this is k-mer size for deBruijn graph values between 25 and 127 are supported, auto will compute the optimal size based on the read data and GC content
GRAPH_KMER_SIZE = auto

#set this to 1 for all Illumina-only assemblies
#set this to 0 if you have more than 15x coverage by long reads (Pacbio or Nanopore) or any other long reads/mate pairs (Illumina MP, Sanger, 454, etc)
USE_LINKING_MATES=1

#specifies whether to run mega-reads correction on the grid
USE_GRID=0

#number of cpus to use, set this to the number of CPUs/threads per node you will be using
NUM_THREADS = 16

#this is mandatory jellyfish hash size -- a safe value is estimated_genome_size*20
JF_SIZE = 200000000

#ILLUMINA ONLY. Set this to 1 to use SOAPdenovo contigging/scaffolding module.  
#Assembly will be worse but will run faster. Useful for very large (>=8Gbp) genomes from Illumina-only data

SOAP_ASSEMBLY=0

END 
